# GIGYF2 mediates post-transcriptional mRNA repression through recruitment of the CCR4/NOT complex

**DOI:** 10.1101/181776

**Authors:** Cinthia Claudia Amaya Ramirez, Petra Hubbe, Nicolas Mandel, Julien Béthune

## Abstract

Initially identified as a factor involved in tyrosine kinase receptor signalling, GRB10-interacting GYF protein 2 (GIGYF2) has later been shown to interact with the 5’ cap-binding protein m4EHP as part of a translation repression complex, and to mediate post-transcriptional repression of tethered reporter mRNAs. We recently observed that GIGYF2 also interacts with the miRNA-induced silencing complex and modulates its translation repression activity. Here we have further investigated how GIGYF2 represses mRNA function. In RNA tethering reporter assays we show that GIGYF2 exerts its action through a combination of translational repression and stimulated mRNA decay. Using truncation variants we identify two distinct effector domains within GIGYF2. In this assay GIGYF2-mediated repression is independent of m4EHP but dependent on the deadenylation activity of the CCR4/NOT complex. We further show that GIGYF2 interacts with multiple subunits of the CCR4/NOT complex and interestingly depletion of the CNOT1 scaffold subunit does not affect GIGYF2-mediated repression. Finally, we identify endogenous mRNA targets of GIGYF2 that recapitulate m4EHP - independent repression. Altogether, we propose that GIGYF2 has two distinct mechanisms of repression: one depends on m4EHP binding and affects translation, the other is m4EHP-independent and relies on the deadenylation activity of the CCR4/NOT complex.

## INTRODUCTION

Recent mRNA interactome studies demonstrated that the repertoire of eukaryotic RBPs is much larger than previously anticipated, including numerous proteins with non-canonical RNA binding domains (1-3). Moreover it is clear that the final regulation of a given mRNA results from the combinatorial action of several RBPs, defining a ribonucleoprotein complex (RNP) that can remodel according to cellular context and external signaling cues (4). Hence it is important to identify and characterize RBPs in order to fully understand how genes are post-transcriptionally regulated. Argonaute proteins are the core component of the microRNA-induced silencing complex (miRISC) that uses the small non-coding microRNAs (miRNA) as guides to find mRNA targets through partial sequence complementarity (5). In mammalian cells, miRISC silences its targets through an initial translation repression step, followed by stimulated deadenylation and decay (6). In order to find additional factors that modulate the activity of the miRISC, we recently described a novel split-BioID conditional proteomics approach that allowed us to specifically probe Ago2-interacting proteins in the context of miRNA-mediated repression (7). One of the major hits we found was Grb10-interacting GYF protein 2 (GIGYF2). GIGYF2 was initially identified as a ubiquitously expressed murine Grb10 - interacting protein involved in IGF-I and insulin receptor signaling regulation (8). GIGYF2 has a potential role in the development of Parkinson’s disease, though this is under debate due to contradictory evidence from different reports (9,10). Knockout studies in mouse are supportive of a role of GIGYF2 in the development of neurological disorders. Indeed, while homozygous animals died shortly after birth due to their inability to feed, heterozygous mice appeared to develop normally but, with age, gradually showed motor dysfunction correlated with signs of neurodegeneration and defect in IGF signaling (11). Interestingly, GIGYF2 was identified as part of a translation repression complex when bound to the mammalian cap binding protein m4EHP (12). In this complex, GIGYF2 is proposed to serve as a bridging protein between RBPs and m4EHP to promote translational repression. Recently, the RBP tristetraprolin (TTP) was shown to partly rely on this mechanism to repress its target mRNAs (13). Supporting a potential role of GIGYF2 in the post-transcriptional regulation of mRNAs, a previous study described an interaction of GIGYF2’s isolated GYF domain with components of stress granules including miRISC components (14). Moreover, in an RNA-tethering assay, GIGYF2 was shown to repress mRNA function (15). In a preliminary characterization, we have shown that GIGYF2 directly but transiently interacts with the miRISC component GW182 (also known as TNRC6 in mammals) (7). The interaction is mediated by a conserved PPGL motif within GW182 that is recognized by the GYF domain of GIGYF2. Importantly, we also showed that GIGYF2 positively regulates the activity of the miRISC at early stages of miRNA-mediated silencing (7) when translation repression is the dominant mechanism of repression. In this study we set out to further characterize GIGYF2. We report that GIGYF2 is a repressor of mRNA function, independent of the previously reported interaction with m4EHP. Moreover we show that GIGYF2 is an RBP that can directly bind to mRNAs. We mapped two effector domains within GIGYF2 and show that its repressing activity is dependent on the deadenylation activity of the CCR4/NOT complex. We further provide evidence that GIGYF2 interacts with the CCR4/NOT complex through interfaces that involve multiple CNOT subunits. Importantly, using microarray analysis and RNA-IP we identify endogenous targets of GIGYF2-mediated silencing that are repressed in an m4EHP-independent manner. Altogether, our data unravel a previously overlooked function of GIGYF2 as an RBP that represses its target mRNAs by stimulating their decay independent of the previously described interaction with m4EHP.

## MATERIAL AND METHODS

### Plasmids

Plasmids for the tethering reporter assay pRL-5boxB, pGL3-FL, pCIneo-λHA-LacZ (16), pCIneo-λHA - and HA-TNRC6C (17) were a kind gift from Witold Filipowicz. The plasmid encoding eGFP-GIGYF2 (14) was a kind gift of Christian Freund (Freie Universität Berlin). The plasmids encoding CNOT7, CNOT8 and CNOT9 were a kind gift of Sebastiaan Winkler (University of Nottingham), the plasmids containing CNOT1, CNOT2 and CNOT3 cDNA were a gift from Elisa Izaurralde (Addgene 37370, 37371 and 37372) and the plasmid encoding CNOT6 was a kind gift from Ann-Bin Shyu (University of Texas). Plasmids encoding dominant negative mutants of Caf1 and CCR4a in a pBEG-3xFLAG vector were obtained from Marina Chekulaeva (Max Delbrück Center for Molecular Medicine, Berlin). Plasmids encoding λHA-GIGYF2, and its deletion fragments were generated by PCR amplification of the corresponding fragments, digested with MfeI and NotI, and cloning into EcoRI and NotI - digested pCIneo-λHA vectors. For the generation of pCIneo-HA-LacZ and - HA-GIGYF2, the respective λHA plasmids were digested with NheI and XhoI, the overhangs filled and the plasmid re-ligated. To generate pCIneo-λHA-mutGIGYF2, previously described mutations (12) were introduced in pCIneo - λHA-GIGYF2 by site directed mutagenesis. pMIR-FL-5boxB was generated by exchange the 3’UTR of pMIR - HMGA2 3’UTR (18) with a 5boxB PCR fragment using SacI and NaeI. The plasmids encoding CNOT subunits or Ago2 fused with a GFP tag, were cloned by PCR amplification of the corresponding ORF, and ligated into peGFP vector using BamHI and XhoI. Removing GIGYF2 from peGFP-GIGYF2 using BamHI and XhoI generated the plasmid peGFP-C3, then the overhangs were filled and the plasmid was re-ligated. The cDNA from m4EHP was obtained using reverse transcriptase PCR using total RNA from HeLa 11ht cells and cloned into EcoRI/NotI-digested pCIneo vector that contains an HA tag or in an SbfI/NotI-digested pBEG-3xFLAG vector. Correctness of all plasmids was verified by Sanger sequencing.

### Cell Culture, transfections and RNAi

Hela 11ht cells (19) were grown in DMEM medium (SIGMA) supplemented with 2mM L-glutamine (GIBCO), 10% (v/v) tet-free FBS (TH. Geyer) and 200µg/mL G418 (Sigma). Cells were regularly tested for mycoplasma contamination.

DNA transfections were performed using Polyethylenimide (PEI) (Polysciences, Inc) in a ratio 1:2 (DNA: PEI). Cells had their medium changed directly before transfection. Plasmid DNA (3 µg for 6 well and 10-15 µg for 10 cm), PEI and serum-free DMEM were mixed, incubated for 10 min at RT and added to the cells. In tethering experiments, cells were transfected with 2ng pRL-5boxB, 50ng of pGL3-FL and 10ng HA or λHA fusion constructs per well or alternatively with 25ng pMir-FL-5boxB and 20ng HA or λHA fusion constructs per well of a 96 well plate. When indicated 69ng of each plasmid encoding CNOT6 and CNOT7 catalytic mutants were co-transfected. For experiments in which the same lysate was used for RNA extraction, Western blot and tethering assay, the transfection was performed either using 200ng pMIR-FL-5boxB and 200 ng HA-λHA fusion constructs per well of a 6 well plate or 25ng RL-5boxB, 300 ng of pGL3-FL and 300ng HA or λHA fusion constructs. For mRNA half-lives measurements, cells were seeded in a 12 well plate, transfected with tethering reporters 24h after seeding, and after an additional 24h treated with 5µg/ml Actinomycin D (Sigma) as indicated.

siRNAs and esiRNAs were previously described (7). Transfections of siRNA/esiRNA or cotransfections with DNA were performed with the jetPRIME reagent (Polyplus Transfection). siRNAs/esiRNAs were transfected at a final concentration of 50nM following the manufacturer’s instructions. The medium of the cells was changed 24h after transfection and the cell were lysed after 48h.

### Luciferase assay

Cells were lysed 24h after transfection using passive lysis buffer (Promega). In case the same lysate was used for RNA extraction, cells were lysed using a cytoplasmic RNA lysis buffer (50 mM Tris-HCl pH 7.4, 150 mM NaCl, 1mM EDTA pH 8, 0.5% NP-40) for 20 min at 4°C. After centrifugation at 10,000 *g* for 10 min at 4°C, part of the supernatant was used for a Luciferase Assay, the equivalent of 10-15 µg protein content was kept for a Western blot analysis and the rest was used for RNA extraction. Firefly (FL) and Renilla luciferase (RL) activities were measured with a Xenius XL microplate luminometer (SAFAS, Monaco) using the Dual-Luciferase Reporter Assay System (Promega).

### Immunofluorescence

Immunofluorescence was performed using HeLa-11ht cells seeded in a 12 well removable chamber slide (Ibidi). Cells were fixed with 3.7% Formaldehyde for 10 min at RT followed by three washing steps with PBS, each 3 min. Permeabilization was performed with 100% ice-cold methanol for 5 min at RT, followed by three washing steps with PBS, each 3 min. Before incubation of primary antibodies for 1h at RT, cells were blocked in 3%FBS in PBS for 30min at RT. Cells were washed 3 times with PBS for 3 min each wash and then incubated with secondary antibodies for 1h at RT followed by three washing steps in PBS, each 3 min. DAPI staining (300nM) was done for 5 min at RT and cells were afterward washed again two times with PBS prior fixation with Fluoromount-G (Southern Biotech). Pictures were taken with the 63x objective of a Nikon Ni-E widefield microscope controlled by the Velocity software at the Heidelberg Nikon imaging center (Heidelberg). Minimum and maximum displayed values can be found in Supplementary Table S1. Cutoffs values were set against cells, which had not been treated with primary antibodies.

### Antibodies

For Western blot analysis, antibodies directed against the following proteins were used: GFP (SC-9996, Santa Cruz, 1:2000), α-tubulin (T6074, Sigma, 1:10000), α-tubulin (Ab18251, Abcam 1:10000), GAPDH (60004-1lg, Proteintech, 1:20000), HA (16B12, Covance, 1:1000), GIGYF2 (NBP2-12812, Novusbio, 1:3000), FLAG-tag (F1804, Sigma, 1:500). Secondary antibodies were: IRDye 800CW goat anti-rabbit (LICOR, 1:10000), IRDye 680RD goat anti-rabbit (LICOR, 1:10000), IRDye 800CW goat anti-mouse (LICOR, 1:10000) and Alexa 680 goat anti-mouse (A21057, Life Technologies 1:10000). For Immunoprecipitation: HA (Covance), GIGYF2 (Novusbio) and mouse or rabbit IgG (Sigma). For Immunofluorescence: GIGYF2 (NBP2-12812, Novusbio, 1:500), HuR (sc-5261, Santa Cruz, 1:50), p70 S6 Kinaseα (nuclear) that strongly crossreacts with Hedls (20) a marker of cytoplasmic P-bodies (sc-8416, Santa Cruz, 1:500), Calnexin (610523, BD Transduction Laboratories, 1:100), anti-mouse Alexa 488 (A28175, Invitrogen, 1:1000) and anti-rabbit Alexa 633 (A21070, Invitrogen, 1:1000).

### GFP pull-down and Immunoprecipitation

For GFP pull-down assays, HeLa 11ht cells grown in a 10-cm dish were transfected with 7.5 μg plasmid expressing eGFP-CNOT fusion proteins and 7.5 μg plasmid expressing HA-GIGYF2 or its deletions fragments. 24h after transfection, cells were lysed in 1.4 mL lysis buffer A (50 mM Tris-HCl pH 7.4, 150 mM NaCl, 2 mM EDTA pH 8, 0.5% NP-40, 0.5 mM DTT, 1 x Complete protease inhibitor (Roche) for 30min at 4°C, and cleared lysates were then treated with micrococcal nuclease (NEB, 14 gel Unit/µl) and 2.5 mM CaCl_2_ for 25 min at RT, we have tested that this treatment eliminates RNA - dependent interactions. The lysates were then incubated with 25 µl pre-equilibrated GFP-Trap magnetic beads (Chromotek) for 2-3h at 4°C on a rotating wheel. Beads were then washed three times with the same equilibration buffer 1 (50mM Tris pH 7.4, 300 mM NaCl, 1 mM MgCl2, 0.1% NP-40) and GFP-fusion proteins were eluted by boiling the beads at 95°C for 5 min in Laemmli SDS loading buffer.

For immunoprecipitation assays, Hela 11ht cells were grown in a 10-cm dish were transfected with 10 μg plasmid expressing HA-m4EHP, HA-GIGYF2, or variants thereof (HA-IP), or with 7.5 µg plasmid for FLAG-m4EHP expression and 7.5 µg plasmid coding for HA-LacZ, HA-GIGYF2 or its mutant (FLAG-IP). 24h after transfection, cells were lysed as in the GFP pulldown assay. The lysates were then incubated with 25 µl pre-equilibrated protein G magnetic beads (NEB) previously coupled with 5 μg of the relevant antibody or control IgG overnight at 4°C on a rotating wheel. Beads were then treated as for the GFP pulldown assay, using an equilibration buffer 2 (50mM Tris pH 7.4, 300 mM NaCl, 1 mM MgCl2, 0.5% NP-40). For FLAG Immunoprecipitations, bound proteins were eluted with 150 ng/µl FLAG Peptide (BACHEM) in TBS for 1h at 4°C.

### RNA-Immunoprecipitation

25 µl of protein G magnetic beads (NEB) were washed with equilibration buffer 2 and coupled with 5 μg of anti-GIGYF2 or rabbit IgG overnight at 4°C on a rotating wheel in equilibration buffer 2 supplemented with 35 μg Heparin, 50 μg tRNA from *E. coli* and 50U/mL Ribolock. On the next day, two confluent 15-cm dishes of HeLa-11ht cells were used. After washing twice with ice-cold PBS, cells were scrapped in PBS and harvested by centrifugation at 300 *g* for 10 min at 4°C. Cells were lysed in 1.2 mL lysis buffer A, supplemented with 50U/mL Ribolock (Thermo Scientific) for 30 min on ice and the lysates cleared by centrifugation at 10,000g x 15 min at 4°C. The cleared lysates were then divided and incubated with the coupled anti-GIGYF2 or control beads for 3h at 4°C on a rotating wheel. The beads were then washed three times with equilibration buffer 2 supplemented with 1 x complete protease inhibitor and 50U/mL Ribolock. For elution the beads were re-suspended in Proteinase K Buffer (300 mM NaCl, 200 mM Tris-HCl pH 7.5, 25 mM EDTA pH 8, 2% SDS) with 200 μg of Proteinase K (NEB) and incubated at 65°C for 15min.

### RNA extraction, cDNA synthesis and qPCR

For RNA extraction and purification, cells or RNA-IPed material were mixed with TRI or TRI LS Reagent (Sigma) and purified on affinity columns using the Direct-Zol RNA miniprep kit (Zymo) that includes a DNase-treatment step. For cDNA synthesis, 1 μg of purified RNA for first treated with DNaseI (NEB) and then reverse-transcribed using the Transcriptor first strand cDNA synthesis kit (Roche) with oligo(dT) or random hexamers primers. For qPCR analysis, the cDNA were typically diluted in water (1:5) and used as a template for SYBR Green based qPCR using the SYBR Green qPCR Mix (Roche) and gene specific primers (See Supplementary Table S2) in a Step One Plus Real-Time PCR System (Applied Biosystems). Each reaction was performed in technical duplicates or triplicates. Relative expression levels were calculated with the ΔCt and the data was normalized to GAPDH. Relative expression levels were calculated with the formula 2^-(ΔCt)^, where ΔCt is Ct(RL or FL)-Ct(GAPDH) and Ct is the equivalent cycle number at which the chosen threshold is crossed.

### RNA binding assay

The RNA binding assay was adapted from a previous report (21). HeLa 11ht cells grown in a 10 cm dish were transfected with 10 μg plasmid expressing eGFP, eGFP-Ago2 or eGFP-GIGYF2 fusion proteins. 24h after transfection, cells were washed once with PBS and 4mL of PBS was added to the dishes. Cells were then placed on ice and immediately irradiated with 0.15J/cm^2^ UV light at 254nm (UV Stratalinker 1800, Stratagene). Cells were scrapped, collected by centrifugation and resuspended in 300 µl lysis buffer B (100mM KCl, 5mM MgCl2, 10 mM Tris pH 7.4, 1% NP-40, 1mM DTT, 100 units/mL Ribolock, 1X protease inhibitor cocktail and 200 µM ribonucleoside vanvadyl complex (NEB)). Lysates were incubated on ice for 10 min, snap-frozen, and thawed to achieve complete lysis. Lysates were centrifuged for 10min at 10 000 rpm at 4°C and the supernatant was dispatched into three tubes. Samples were mixed with 400 µl of dilution buffer (500 mM NaCl, 1 mM Mg2Cl, 0.05% SDS, 0.05% NP-40, 50 mM Tris-HCl pH 7.4, 100 units/mL Ribolock and 1X protease inhibitor cocktail) and 10 µl of pre-equilibrated GFP-Trap A beads (Chromotek). GFP beads were preequilibrated as in GFP pulldown experiments (see above). GFP-tagged proteins were immunoprecipitated for 2h at 4° on a rotating wheel. Beads were then washed with 500µl medium salt buffer (250 mM NaCl, 1 mM Mg2Cl, 0.025% SDS, 0.05% NP-40, 20 mM Tris-HCl pH 7.4, 50 units/mL Ribolock, 1X protease inhibitor cocktail) and incubated for 15 min at 4°C on a rotating wheel with 250 µl of blocking solution (200mM LiCl, 20mM Tris-HCl pH 7.5, 1 mM EDTA pH 8, 0.01% NP-40, 100 µg/mL *Escherichia Coli* tRNA, 100 µg/mL BSA, 50 units/mL Ribolock, 1X protease inhibitor cocktail). Beads were then incubated for 1h at 4°C on a rotating wheel with 250 µl of hybridization buffer (500 mM LiCl, 20 mM Tris-HCl pH 7.4, 0.05% LiDS, 1 mM EDTA pH 8, 5 mM DTT, 0.01% NP-40, 8nM oligo(DT)25 WellRED (Sigma), 100 units/mL Ribolock and 1X protease inhibitor cocktail) and the excess of fluorescent was removed by washing once with 500 µl of washing buffer A (500 mM LiCl, 20 mM Tris-HCl 7.4, 0.01% LiDS, 0.01% NP-40, 1 mM EDTA pH 8, 5 mM DTT, 50 units/mL Ribolock, 1X protease inhibitor cocktail) and twice with 500 µl of washing buffer B (200 mM LiCl, 20 mM Tris - HCl 7.4, 0.01% LiDS, 0.01% NP-40, 1mM EDTA pH 8, 5 mM DTT, 50 units/mL Ribolock, 1X protease inhibitor cocktail). Beads were re-suspended in 100 µl of washing buffer B and transferred to a 96-well optical plate (UV Plates, 4titude).

### Fluorescence measurements

Fluorescence signals from the RNA binding assay were measured using a Spectra Max M5 plate reader (Molecular Devices), using the following parameters: eGFP: excitation 475 nm, emission 509 nm; WellRED: excitation 650nm, emission 670 nm. The measurements were done from the bottom of the plate.

### Microarray Analysis

RNA extracted from Hela 11ht WT cells and cells overexpressing GIGYF2 from three biological replicates was sent to the Genomics Core Facility (EMBL, Heidelberg) for Affymetrix GeneChip array analysis. RNA samples (500ng) were processed and labeled for array hybridization using the Ambion WT Expression kit (Life Technologies, catalogue number 4411974). Labeled, fragmented cDNA (Affymetrix GeneChip^®^ WT Terminal Labeling and Controls Kit, catalogue number 901524) was hybridized to Human Gene 2.0 arrays for 16 hours at 45°C (at 60 rpm) (Affymetrix GeneChip^®^ Hybridization, Wash, and Stain Kit, catalogue number 900720). Arrays were washed and stained using the Affymetrix Fluidics Station 450, and scanned using the Hewlett-Packard GeneArray Scanner 3000 7G. Data analysis was performed with the Transcriptome Analysis Console (TAC) Software (Affymetrix).

## RESULTS

### GIGYF2 is part of mRNPs that shuttle to stress granules

We have recently shown using a novel conditional proteomics approach that GIGYF2 is associated with a complex that contains both miRISC core components Ago2 and GW182, and validation by co-immunoprecipitation revealed that the association is stabilized by RNA (7). Using recombinant protein fragments we found that the interaction is mediated by a direct binding involving the GYF domain of GIGYF2 and a conserved PPGL motif of GW182 (7). Moreover, depletion of GIGYF2 alleviated miRNA-mediated translation repression (7). To seek additional evidence for the presence of GIGYF2 in ribonucleoprotein complexes (RNP), we analyzed its cellular localization. In agreement with previous data (14), indirect immunofluorescence showed that in normal conditions endogenous GIGYF2 is mainly localized in the cytoplasm with extensive co-staining with the endoplasmic reticulum marker calnexin (Figure 1A). By contrast, upon sodium arsenite-induced cellular stress, GIGYF2 translocates to stress granules (Figure 1B, C) as evidenced by extensive colocalization with HuR, an RBP that serves as a marker for these cytoplasmic RNA granules (22). As in certain conditions HuR is reported to also localize to P-bodies (23), another class of cytoplasmic RNA granules, we also performed co-stainings with an antibody crossreacting with Hedls a marker of P - body (24). Neither in normal nor in stress conditions (Figure 1D, E) could we observe significant colocalization. Altogether, this suggests that GIGYF2 is part of RNPs that can specifically localize to stress granules.

**Figure 1.**
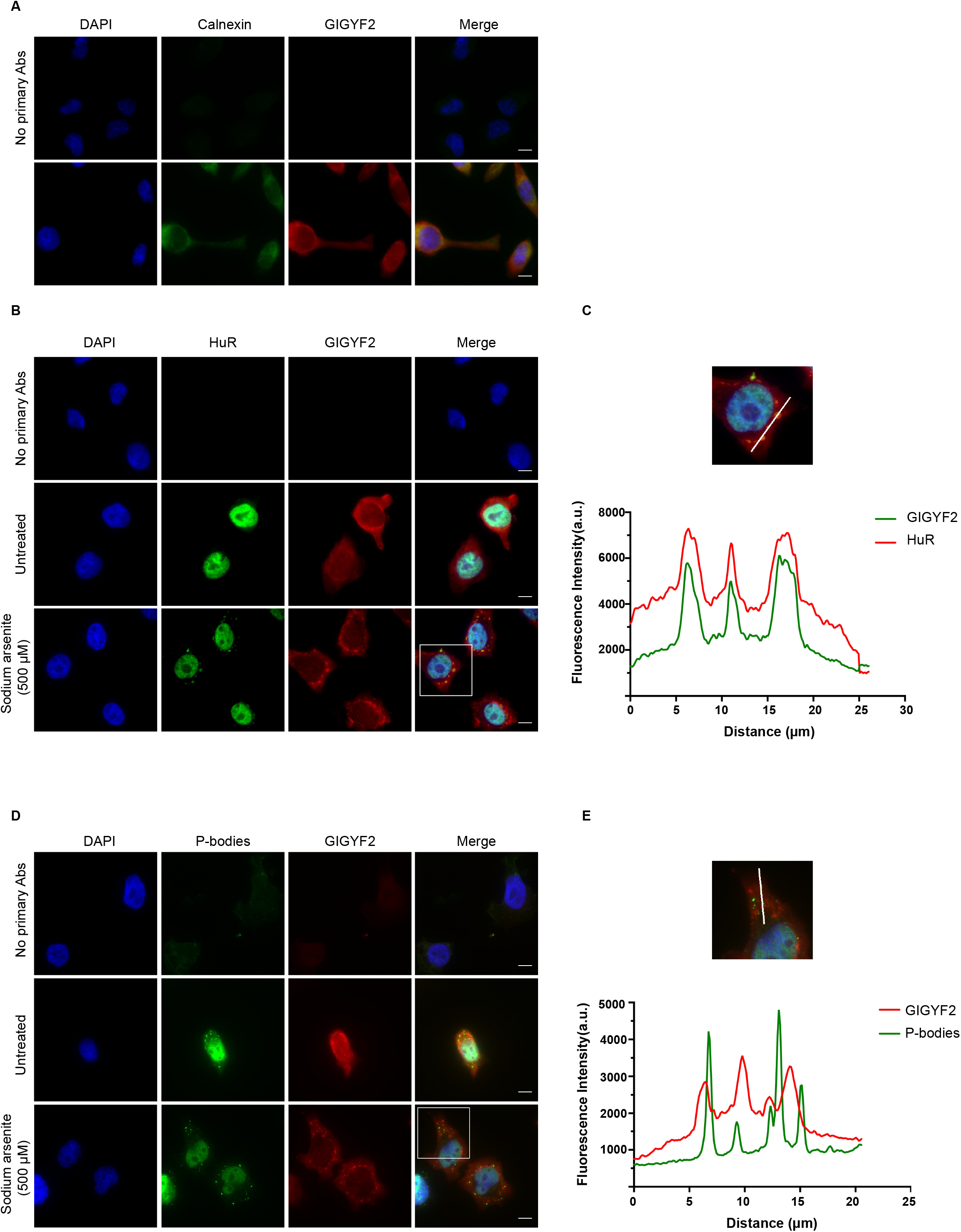
GIGYF2 is part of RNPs. (A) Immunofluorescence of HeLa-11ht cells decorated with antibodies against GIGYF2 and calnexin (ER marker) as indicated. (B) Immunofluorescence of HeLa cells decorated with antibodies against GIGYF2 and HuR (stress granules marker). When indicated, cells were pre-treated with 500µM Sodium Arsenite for 30 min. (C) Enlargement of the area of interest shown in the merged image of (B) and fluorescence intensity profile [arbitrary units (a.u.)] across the marked white line. (D) Same as (B) with antibodies against GIGYF2 and pp70 s6 kinase (that cross-reacts with Hedls a marker of P-bodies). (E) Same as (C) for the area indicated in (D). Scale bar 10µm. Representative images of three independent experiments are shown.

### GIGYF2 represses mRNA function

Since GIGYF2 is part of RNPs, we asked if it modulates mRNA function as was suggested before (15). To this end, we used an RNA-protein tethering assay in HeLa cells (Figure 2A). GIGYF2 fused to a λN peptide and an HA-tag (λHA-GIGYF2) was co-expressed with a thymidine kinase (TK) promoter-driven firefly luciferase mRNA reporter (termed FL-5boxB) containing, in its 3’UTR, five B-box hairpins that are specifically bound by the λN peptide (25). As a normalization control, a Renilla luciferase reporter (termed RL) without B-box was co-expressed with the FL-5boxB reporter. As protein controls, we used a λN peptide fused to HA-tagged β-galactosidase (λHA-lacZ), which is not expected to repress FL-5boxB, and a λHA fusion to the GW182 protein TNRC6C (λHA-TNRC6C), which is known to very efficiently repress boxB-tagged mRNA reporters in tethering assays (17). In addition, HA-tagged GIGYF2, LacZ and TNRC6C were used as additional controls to ensure that any observed effect was due to the tethering of the proteins to the FL-5boxB reporter mRNA and not merely to pleiotropic overexpression effects. To calculate the repression induced by the tethered proteins, the normalized expression of the FL-5boxB reporter in the presence of the λHA-tagged proteins was divided by the normalized expression of the FL-5boxB reporter in the presence of the HA-tagged proteins. As expected, λHA-TNRC6C very efficiently repressed the expression of FL-5boxB to 25% of control (Figure 2B). Similarly, λHA-GIGYF2 also significantly repressed luciferase activity of FL-5boxB albeit to a lesser extent than λHA-TNRC6C (to 50% of control). By contrast, no change of normalized FL-5boxB expression was observed on co-expression of λHA-LacZ. Western blot analysis revealed that all the fusion proteins were expressed at similar levels (Figure 2C). Importantly, similar to the TK promoter-driven FL-5boxB reporter, a cytomegalovirus (CMV) promoter-driven RL-5boxB mRNA reporter (with five B-box hairpins in its 3’UTR) was also repressed by λHA - GIGYF2 but not HA-GIGYF2 (Supplementary Figure S1). Hence the observed λHA-GIGYF2-mediated inhibition was not dependent on a specific reporter or promoter. Total RNA was extracted from the samples used in Figure 2 to evaluate if the observed decrease of luciferase activities were correlated to the expression levels of the mRNA reporters. As previously described (17), FL-5boxB mRNA levels only partially correlate with decreased luciferase activity mediated by λHA-TNRC6C (Figure 2B) indicating a repression mediated by both mRNA decay and translational repression. Similarly, decrease in mRNA levels did not match luciferase activities upon expression of λHA-GIGYF2, suggesting that GIGYF2-mediated repression is also due to a combination of mRNA destabilization and translational repression. As similar data were obtained in another study performed in HEK-293 cells (15), this suggests that the mechanism GIGYF2-mediated repression in conserved in different human cell lines.

**Figure 2.**
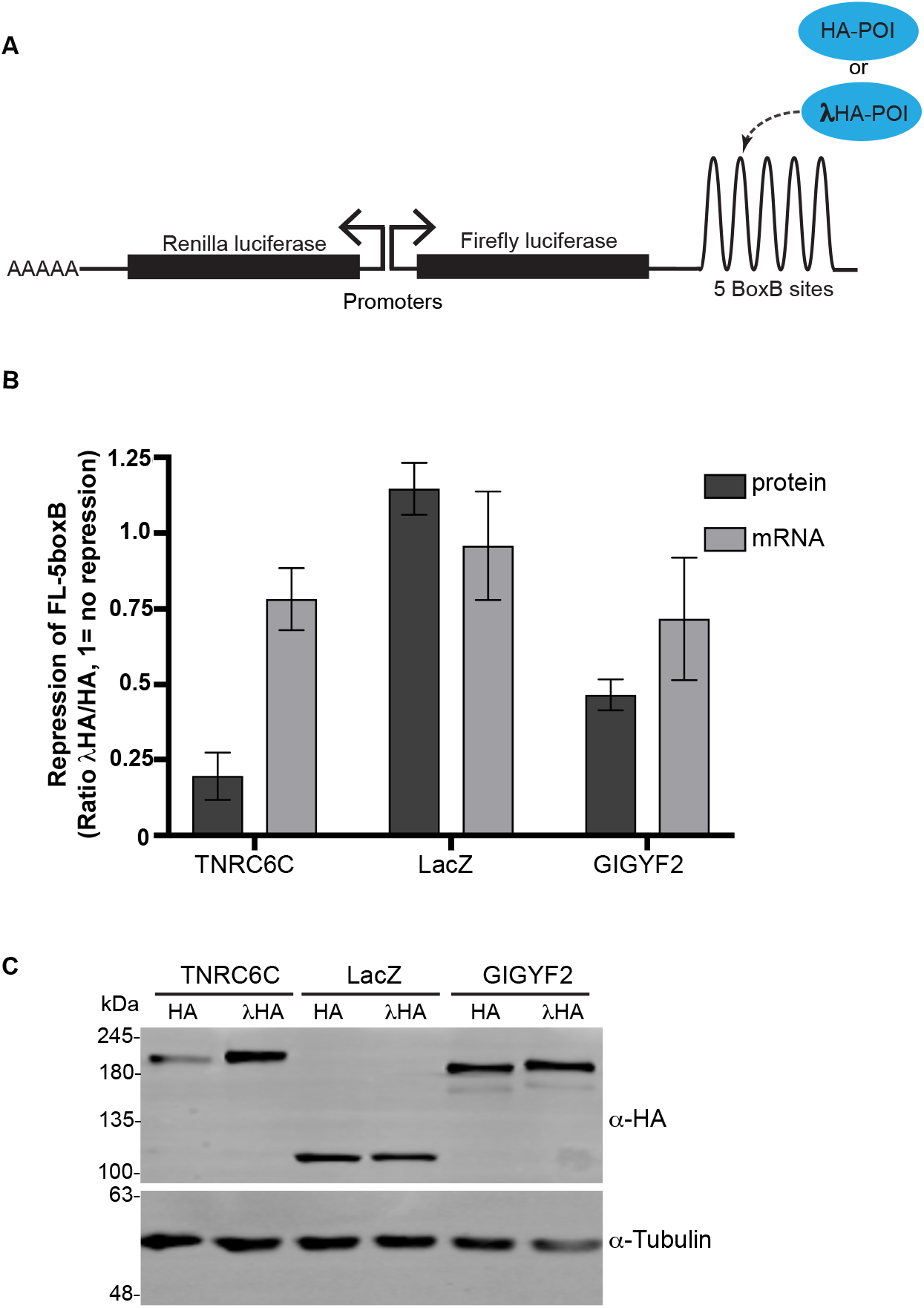
GIGYF2 is a repressor of mRNA function. (A) Schematic of the RNA tethering assay, POI: protein of interest. (B) Repression of the FL-5boxB measured with a dual luciferase assay (protein) or RT-qPCR (mRNA). In both cases, FL-boxB readings were normalized to RL values. The ratios of normalized FL-boxB expression in the presence of the indicated λHA (tethered) over HA-tagged (non tethered) protein were then calculated. Error bars are 95% confidence intervals (n=5 biological replicates) calculated on the logarithmic transformed values of the ratios. The corresponding antilogarithmic values are shown. (C) Expression of the fusion proteins analyzed by Western blotting.

### GIGYF2 is an RNA-binding protein

We then wondered if GIGYF2 directly binds to RNA. This is strongly suggested by mRNA interactome studies in which direct interactors of polyadenylated RNA are comprehensively identified by mass spectrometry following UV-mediated crosslinking of proteins to nucleic acids and pulldown on oligo-dT coupled beads. Indeed, using this unbiased proteomics approach, GIGYF2 was identified as an RBP in HEK-293 (1), Huh7 (26) and in mES (3) cell lines. However, GIGYF2 was absent from the RBP dataset obtained from HeLa cells using the same approach (2). Since we used HeLa cells in this study, we have directly analyzed whether GIGYF2 is also an RBP in this cell line using a recently described fluorescence-based assay (21). To this end GFP-tagged GIGYF2 was expressed in HeLa cells and UV-induced crosslinking was applied prior to cell lysis to capture direct RNA/protein interactions in their native cellular environment (Figure 3A). After cell lysis and affinity purification of tagged GIGYF2 on GFP-trap beads, co-purifying polyadenylated RNA was then detected through hybridization with WellRED-labeled oligo-dT probes. Ratios of red over green fluorescence were then compared with the negative (GFP) and the positive (GFP-tagged Ago2) control proteins. As shown on Figure 3B, a clear and significant enrichment of red fluorescent signal was observed for both Ago2 and GIGYF2 when compared to the control GFP sample, indicating that GIGYF2 is also an RBP in HeLa cells.

**Figure 3.**
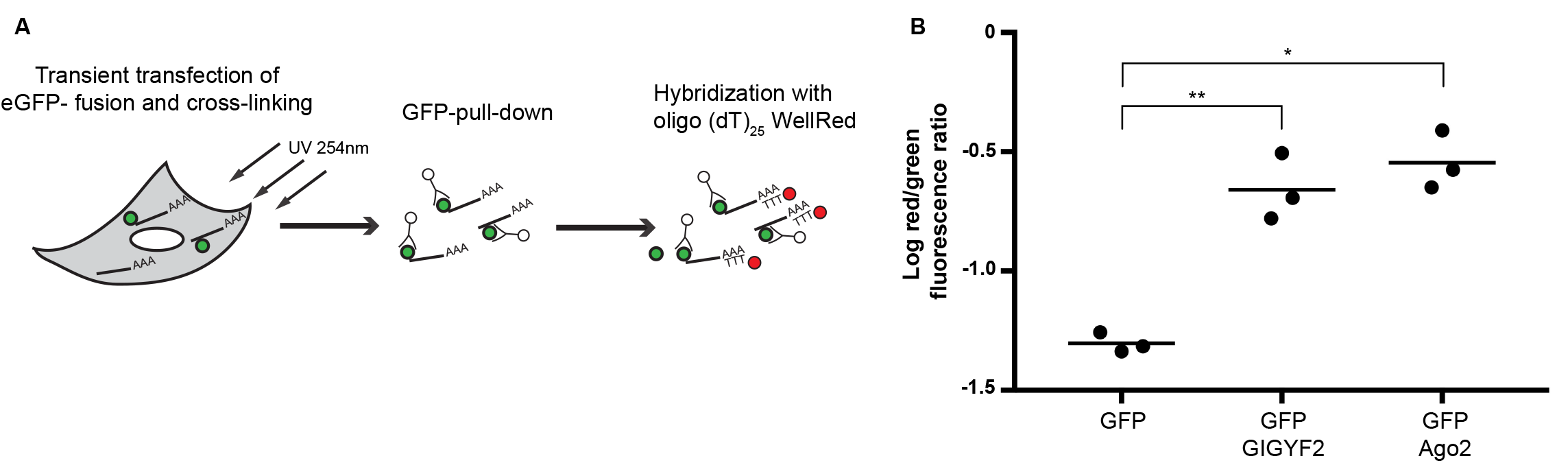
GIGYF2 is an RNA-binding protein. (A) Schematic of the RNA binding assay. (B) HeLa cells transfected with the indicated constructs were irradiated with 254 nm UV light. Lysates were then immunoprecipitated using GFP Trap Agarose beads. Co-immunoprecipitated mRNAs were detected by hybridization with WellRED-labeled oligo(dT)25. Shown are the logarithms of the corresponding red (WellRED) to green (GFP) fluorescence ratios. The graph was generated from three biological replicates each done in three technical replicates. \**P*-value<0.05, \*\**P*<0.01.

### Mapping of GIGYF2 silencing domains

GIGYF2 is a multidomain protein with no clear function assigned to different parts of the protein in the context of mRNA regulation. Using λHA-tagged fusions of truncation variants of GIGYF2, we thus mapped which parts of the protein are responsible for its repressive activity (Figure 4A). Analysis of these variants in the tethering assay revealed that an N-terminal domain (amino acids 1 to 532) of GIGYF2 that comprises the m4EHP-binding motif shows modest repressive activity (Figure 4B, lane 4). This domain is moreover dispensable as a truncation of GIGYF2 lacking the first 532 amino acids (fragment Δ532) shows comparable activity to the full-length protein (Figure 4B, lane 9). When this N-terminal domain was extended to add the GYF domain (fragment 1-596), repressive activity was largely recovered (Figure 4B, lane 5). Extending this latter domain to amino acid 712 led to increased repressive activity but extending further (fragments 1-1025 and 1-1252) did not lead to improved repression. In addition, truncations of GIGYF2 lacking both N-terminal and GYF domains (fragments Δ606 and Δ720) showed repressive activity comparable to the full-length protein. These observations made us hypothesize that GIGYF2 possesses two repressive regions: the GYF domain and an adjacent downstream region. To challenge this hypothesis we expressed two additional fragments of GIGYF2 that encompass the two predicted silencing regions: the isolated GYF domain (fragment 531-596) and a fragment comprising amino acids 607-740 (termed MED for mid effector domain). As both displayed repressive activity comparable to the full-length protein (Figure 4B, last two bars) this strongly suggests that GIGYF2 indeed possesses two distinct silencing domains. Importantly, we verified that all λHA-tagged protein fragments were expressed at comparable levels, which was the case for all except MED that showed much weaker expression levels that did not hampered its silencing activity (Figure 4B). Finally, the interpretation of the RNA-tethering assay data may be compromised in case some of the tested fragments interact with endogenous GIGYF2. To exclude this possibility, various HA-tagged fragments of GIGYF2 were immuno-precipitated from HeLa cell lysates (Figure 4C). By contrast to the positive control HA-m4EHP, no evidence for an interaction with endogenous GIGYF2 could be found for any of the tested GIGYF2 fragments as demonstrated by an absence of signal for endogenous GIGYF2 in the precipitated samples (Figure 4C).

**Figure 4.**
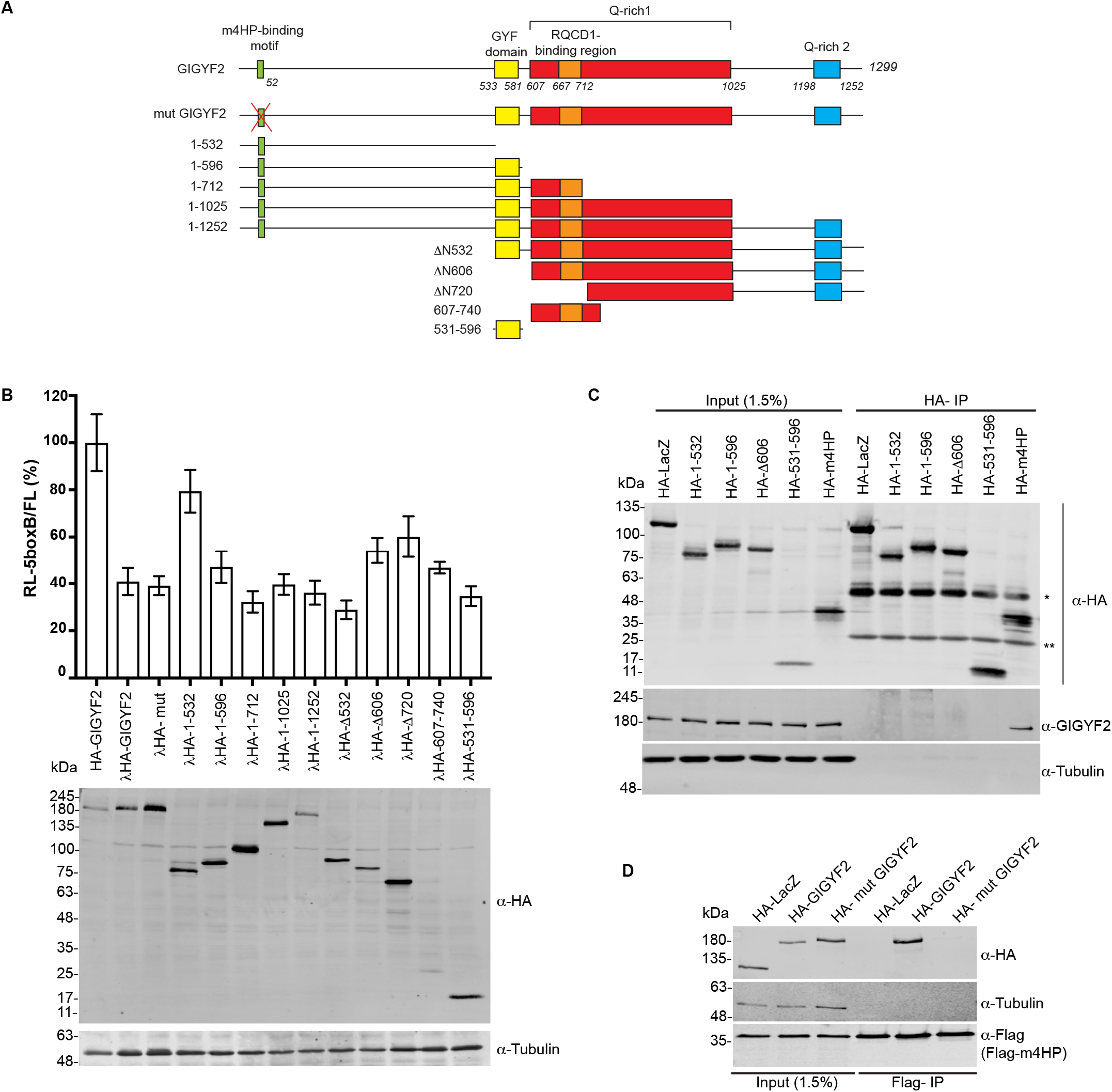
Mapping of GIGYF2 effector domains. (A) Schematic representation of human GIGYF2 and its deletion variants. Positions of binding motif for m4EHP, GYF, Q-rich1, Q-rich2 and RQCD1 - binding domains are indicated. (B) (top) Normalized expression of RL-5boxB measured with a dual luciferase assay. HeLa cells were co-transfected with plasmids expressing the indicated constructs, RL-5BoxB and FL transfection control reporters. RL was normalized to FL and values of normalized RL produced in the presence of HA-GIGYF2 were set to 100%. Mean values are shown with SEM from four to six independent experiments. (bottom) Protein expression was confirmed by Western blotting. (C) Blots of immunoprecipitations performed with an anti-HA antibody from HeLa cells transiently transfected with the indicated constructs, LacZ and m4EHP serve as negative and positive controls respectively. *IgG heavy chain (55 kDa), **IgG light chain (25 kDa). (D) Blots of immunoprecipitations performed with an anti-FLAG antibody from HeLa cells transiently transfected with the indicated constructs.

### GIGYF2-mediated repression is independent of m4EHP

An important observation from the tethering assay was that the N-terminal domain (1-532) that comprises the m4EHP-binding site (12) is not necessary for repression (Figure 4B). As it had been previously proposed that the GIGYF2/m4EHP complex mediates translational repression, we also tested a full-length GIGYF2 variant with a disrupted m4EHP-binding site (mut GIGYF2). In agreement with previous data (12), the introduced mutations efficiently abolished binding of HA-tagged mut GIGYF2 to FLAG-tagged m4EHP in co-IP experiments (Figure 4D). When tested in the tethering assay, λHA-tagged mutGIGYF2 was as effective as the WT protein (Figure 4B, lane 3), demonstrating that GIGYF2 repressing activity is independent of m4EHP.

### GIGYF2 binds to the CCR4/NOT complex

In the context of mammary carcinogenesis, an interaction between GIGYF2 and RQCD1 had been previously described and suggested to enhance association of GIGYF2 with EGFR via the adaptor protein Grb10 (27). The region of GIGYF2 involved in RQCD1 binding had been mapped to amino acids 667-712 (27) and is thus comprised within the MED (Figure 4A). RQCD1 is also known under the name CNOT9 as a subunit of the CCR4/NOT complex that associates with mRNAs and stimulates their deadenylation and decay (28). Since numerous repressors of mRNA function recruit the CCR4/NOT complex to mediate their functions, we asked if GIGYF2 interacts with this complex. To this end, GFP-tagged CNOT subunits were co-expressed with HA-tagged GIGYF2 in HeLa cells and captured on GFP-trap beads (Figure 5). GIGYF2 was found to associate with CNOT1, CNOT9, and the deadenylase CNOT7 (alternatively known as Caf1), and to a lesser extent to CNOT2, CNOT3 and the deadenylase CNOT6 (alternatively known as Ccr4a). As the lysates were pre-treated with the micrococcal nuclease this suggests a binding based on protein-protein interactions.

**Figure 5.**
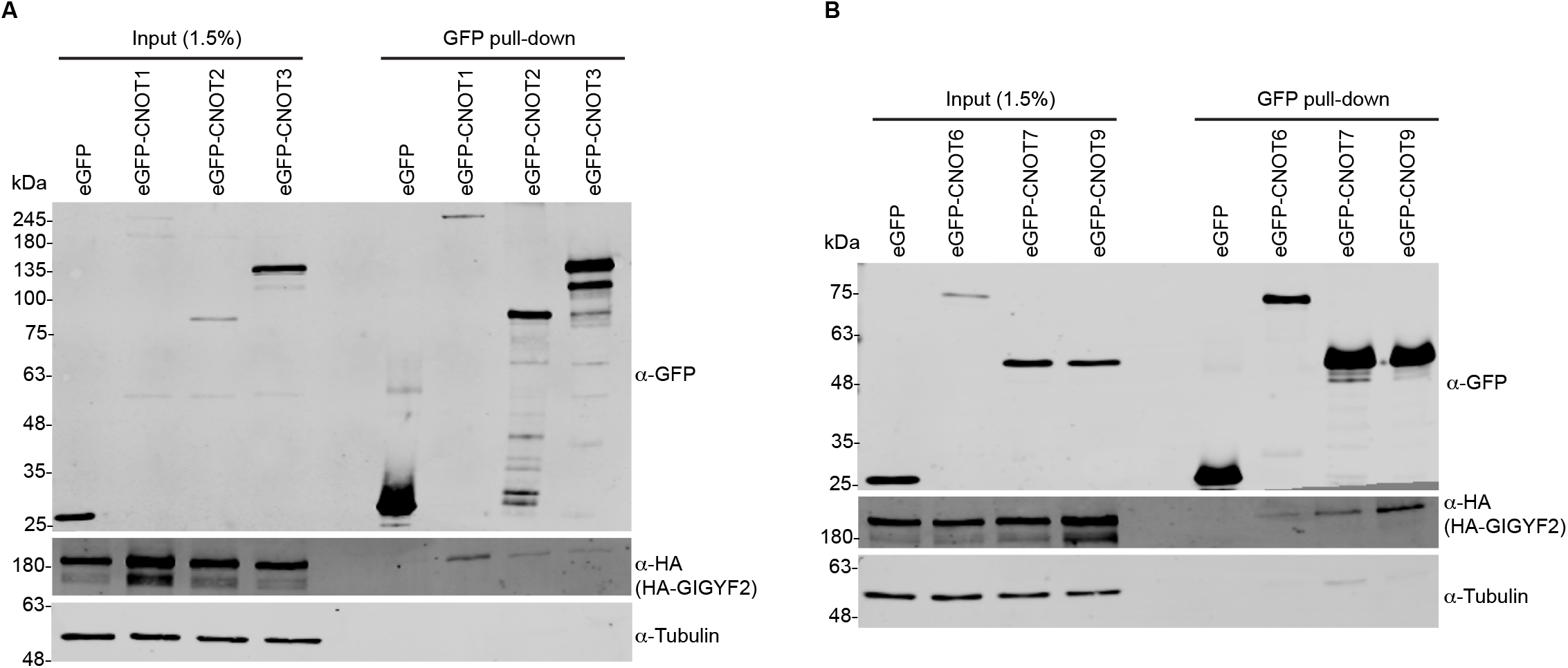
GIGYF2 bind to the CCR4-NOT complex. Blots of GFP pull-down experiments. GFP - fusions of the indicated subunits of the CCR4-NOT complex were co-expressed in HeLa cells with HA-GIGYF2. Cells lysates were immunoprecipitated using GFP-trap magnetic beads. GFP was used as a control. (A) GFP pull-downs with the CNOT1, CNOT2 and CNOT3 subunits. (B) GFP pull-downs with CNOT6, CNOT7 and CNOT9. All the samples were pre-treated with micrococcal nuclease.

### GIGYF2-mediated repression requires the deadenylation activity of the CCR4/NOT complex

The CCR4/NOT complex is a multifunctional complex that is notably involved in many posttranscriptional regulation pathways, including miRNA-mediated silencing (29). We thus wondered whether this complex acts downstream of GIGYF2. While the largest subunit, CNOT1, acts as a scaffold necessary for the assembly of the whole complex, CNOT6 or CNOT6L (alternatively known as Ccr4b) and CNOT7 or CNOT8 (alternatively known as Pop2) are the two subunits with deadenylation activity (28). In a previous study (6), we have shown that a triple knock down of CNOT1, CNOT7 and CNOT8 efficiently disrupts CCR4/NOT-mediated deadenylation. Using the same conditions, GIGYF2-mediated repression was completely alleviated in the tethering assay (Figure 6A). As expected (6,30), the same outcome was observed for TNRC6C-mediated repression (Figure 6B). A previous study showed that CNOT7 and CNOT8 can also lead to translational repression independent of their deadenylation activity (31). To pinpoint which effect is responsible for GIGYF2 - mediated silencing, we used catalytic inactive variants of the two deadenylation subunits CNOT6 (32) and CNOT7 (33). When expressed in cells they act as dominant negative and specifically disrupt CCR4/NOT-mediated deadenylation. In these conditions, GIGYF2-mediated repression was alleviated (Figure 6A), indicating that it strongly relies on mRNA deadenylation. By contrast, TNRC6 was virtually not affected (Figure 6B), reflecting its mechanism of action that involves a strong translation inhibition component in the context of tethering assays (34,35). Interestingly, and by contrast to TNRC6C, GIGYF2-mediated repression was not affected by depletion of the sole CNOT1 subunit (Figure 6A, B). This was unexpected as CNOT1 acts as a scaffold on which other CNOT subunits bind to assemble the full CCR4/NOT complex. This may imply that GIGYF2 can directly bind to deadenylase subunits of the CCR4/NOT complex. Finally, since GIGYF2-mediated repression is dependent on the deadenylation activity of the CCR4/NOT complex, it is expected that GIGYF2 should stimulate the decay rates of mRNAs it is bound to. To verify if this prediction is fulfilled in our tethering assay, boxB-tagged reporter mRNA half-lives were measured by RT-qPCR analysis following an actinomycin D-mediated transcription arrest. As predicted, co-expression of λHA-GIGYF2 led to faster decay kinetics of the boxB-tagged reporter (Figure 6C), but not of a control mRNA (Figure 6D), as compared to conditions in which HA-GIGYF2 was co-expressed. Altogether this suggests that GIGYF2 relies on the deadenylation activity of the CCR4/NOT complex to mediate repression and may interact with the deadenylase subunits independently of CNOT1.

**Figure 6.**
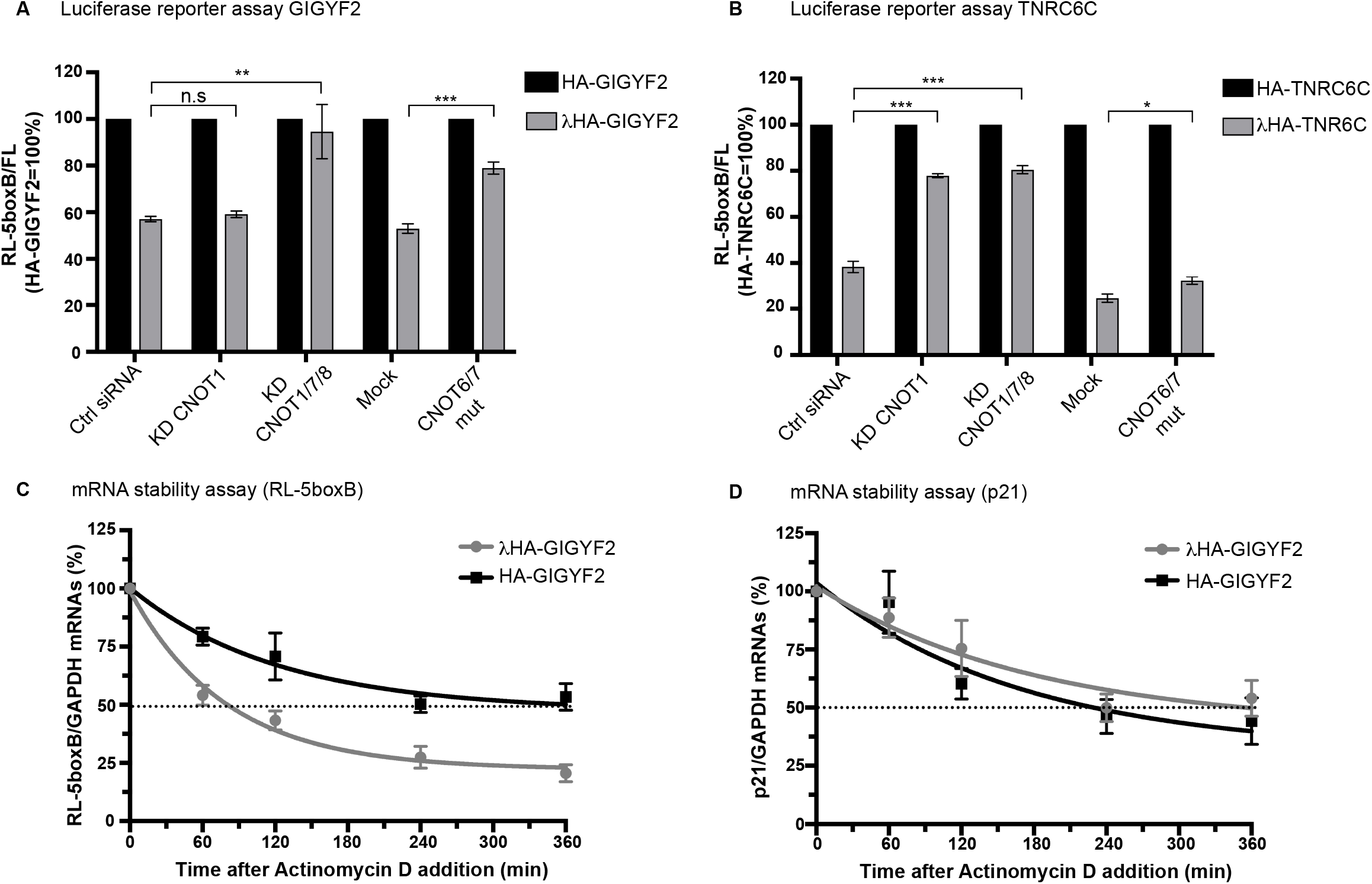
GIGYF2 silencing activity relies on the deadenylation activity of the CCR4-NOT complex. (A) Dual luciferase assay of HeLa cell lysates co-transfected with RL-5boxB, FL and λHA - or HA-GIGYF2 plasmids. When indicated cells were additionally co-transfected with siRNAs against CNOT1, CNOT1/7/8 or a control siRNA, or with plasmids expressing dominant negative variants of CNOT6 and CNOT7 (CNOT6/7 mut). RL readings were normalized to FL and values of normalized RL produced in the presence of the HA-GIGYF2 were set to 100%. Mean values are shown with SEM for three to six independent experiments. (B) Same as (A) with λHA - or HA-TNRC6C plasmids. *P-value<0.05; **p<0.005; ***p<0.001. (C). Expression levels of the RL-5BoxB mRNA reporter over time following Actinomycin D (ActD) mediated transcription arrest in the presence of λHA - or HA-GIGYF2. (D) Same as (C) assessing the p21 mRNA as a control for ActD treatment. RL-5boxB or p21 levels (normalized to GAPDH) were analyzed at the indicated time points by RT-qPCR in five independent experiments and values obtained at time 0 were set to 100%. Error bars are SEM.

### GIGYF2 recruits the CCR4/NOT complex through multiple interfaces

We next wondered which parts of GIGYF2 mediate an interaction with the CCR4/NOT complex. To this end, three truncation variants (1-532, 1-596 and Δ606) used in the tethering assay were coexpressed with GFP-CNOT1/-CNOT7/-CNOT9. The Δ606 variant that comprises the MED was used instead of the MED itself because the latter showed weak expression levels (Figure 4B). After affinity isolation on GFP-trap beads, we could identify the GYF domain as the main CNOT1 and CNOT7 - interacting domain (Figure 7A, B). By contrast CNOT9 preferentially bound to the C-terminal domain of GIGYF2 but also to the GYF domain (Figure 7C). Interestingly the N-terminal fragment (1-532) that hardly showed any activity in the tethering assay was found to bind CNOT1 and CNOT7, suggesting that recruitment of CNOT subunits by this domain is not sufficient to mediate repression. One possible explanation is that this domain is not able to manœuver CNOT subunits in a productive way. Alternatively the (1-532) fragment may not be able to fold correctly on its own. Yet, since the Δ532 truncation variant of GIGYF2 had similar activity than the full-length protein in the tethering assay (Figure 4B), the N-terminal part of the protein does not seem to sensibly contribute to silencing even though it appears to participate in the recruitment of the CCR4/NOT complex. To complement the pulldown studies, tethering assays were performed with the fragments of GIGYF2 upon expression of the dominant negative variants of the CNOT6 and CNOT7 deadenylases. Similar to the full-length protein, all repressive fragments of GIGYF2 showed impaired activity when CCR4/NOT complex - mediated deadenylation was disrupted (Figure 7D). Altogether, the data suggest a physical and functional interaction of GIGYF2 with the CCR4/NOT complex mediated by multiple interfaces involving several domains of GIGYF2 and various CNOT subunits.

**Figure 7.**
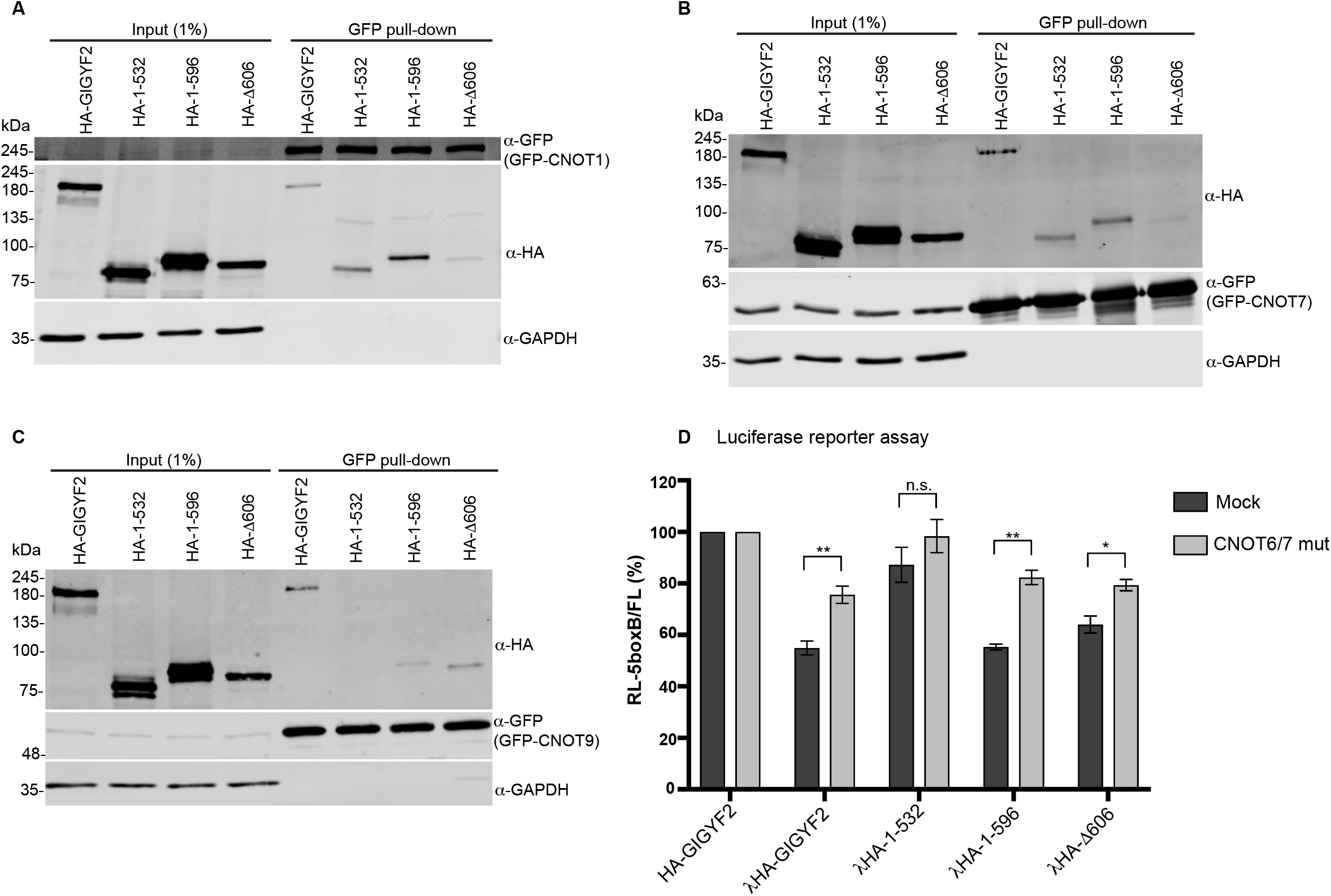
Characterization of the interaction between GIGYF2 and the CCR4/NOT complex. Blots of GFP-pull-down experiments. GFP-fusions of CNOT1 (A), CNOT7 (B) and CNOT9 (C) were co-expressed with the indicated constructs in HeLa cells. Cells lysates were immunoprecipitated using GFP-trap magnetic beads as on Figure 5. (D) Dual luciferase assay of cells transfected with RL - 5boxB, FL and the indicated fusion proteins in the presence (CNOT6/7 mut) or absence (mock) of dominant negative variants of CNOT6 and CNOT7. *P-value<0.01; **p<0.001; n.s., not significant.

### Identification of endogenous targets of GIGYF2

We have shown that, when artificially brought to a reporter mRNA, GIGYF2 stimulates mRNA decay. We next sought to identify potential endogenous RNA targets of GIGYF2 using high-resolution microarrays. To this end, gene expression profiles were compared from WT and GIGYF2 - overexpressing HeLa cells in three biological replicates (Fig 8A). GIGYF2 overexpression led to an upregulation of 132 and a downregulation of 49 transcripts by at least 2-fold (P < 0.05). We then sought to validate six of the downregulated transcripts identified by the microarray approach (AMTN, CA9, SVOPL, COX6B2, LGR5 and COL8A1) and analyzed their expression levels by RT-qPCR using specific primer pairs. All six transcripts showed lower expression upon overexpression of GIGYF2 (Figure 8B). Genuine mRNA targets of GIGYF2 should be up-regulated in conditions where GIGYF2 is depleted. To challenge this prediction, we depleted GIGYF2 from HeLa cells by RNAi and the expression levels of the six selected microarray hits were assessed. We found that four (SVOPL, COX6B2, LGR5, COL8A1) out the six transcripts tested were indeed up regulated upon GIGYF2 depletion (Figure 8C). Finally, to discriminate direct targets of GIGYF2 from transcripts misregulated by secondary effects triggered by non-physiological expression levels of GIGYF2, we perform RNA-IP experiments. Indeed, direct targets of GIGYF2 should associate with the protein. Endogenous GIGYF2 was IPed from HeLa cells and the associated RNA was isolated. From the six selected microarray hits, two transcripts (SVOPL and COL8A1) showed enrichment in the GIGYF2 IP (Figure 8D). Altogether, SVOPL and COL8A1 are very likely *bona fide* direct target of GIGYF2 as they associate with GIGYF2, are down regulated upon overexpression of the protein, and up regulated upon depletion of the protein.

**Figure 8.**
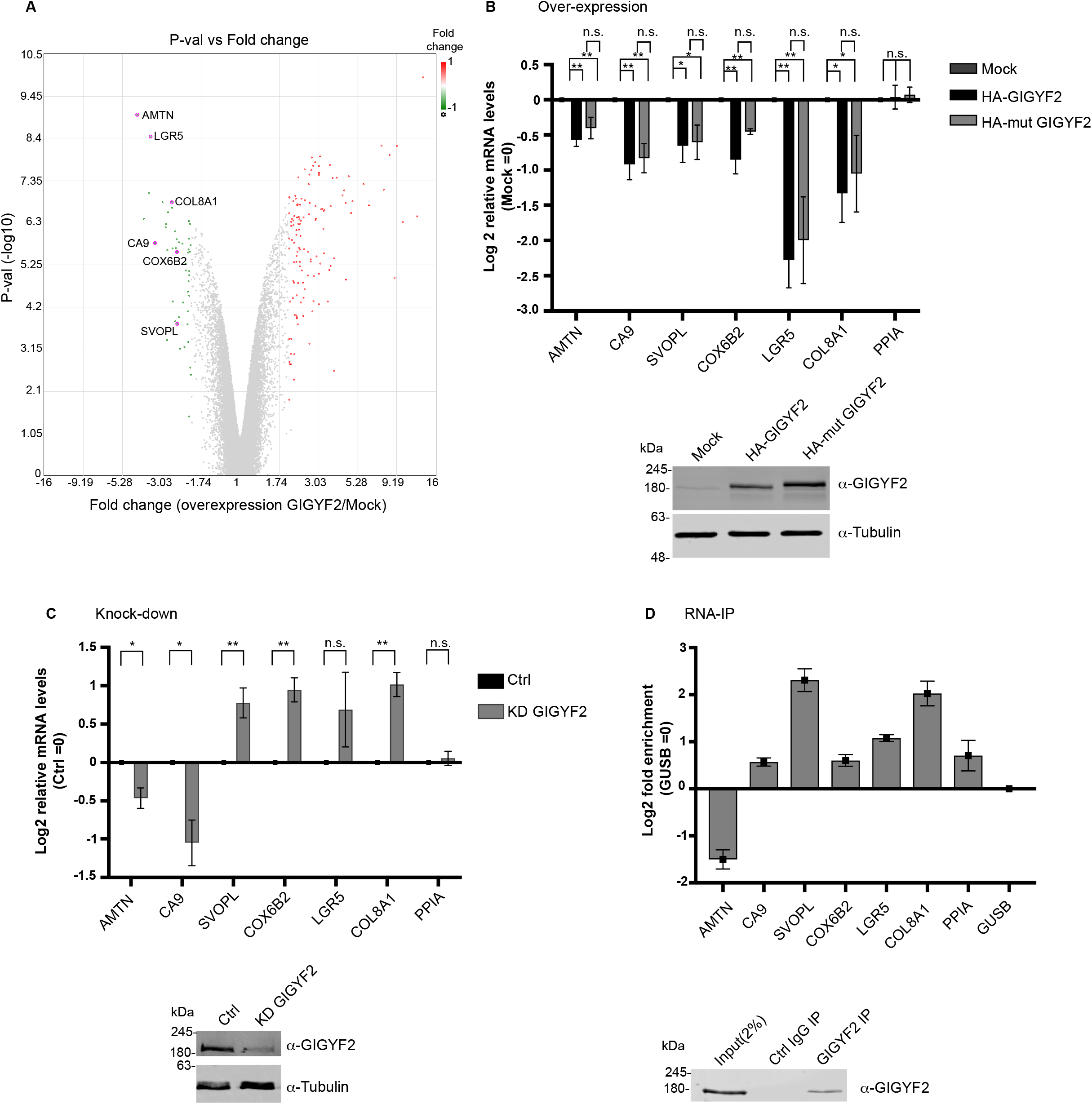
Identification of endogenous targets of GIGYF2. (A) Volcano plot showing mRNA expression level changes upon overexpression of GIGYF2 in HeLa cells analyzed with microarrays. The logarithmic ratios of mRNA levels were plotted against negative logarithmic *P* values of a twosided two samples *t*-test (n=3 biological replicates). Selected transcripts are indicated. (B) Top: expression levels, normalized to GUSB mRNA, of the selected transcripts analyzed by RT-qPCR upon overexpression of GIGYF2 or its m4EHP-binding impaired variant (mut GIGYF2), PPIA is used as a negative control. Log2 values obtained from untransfected cells were set to 0. Mean values are shown with SEM for three to six independent experiments. Bottom: GIGYF2 over-expression was confirmed by Western blotting (C) Top: expression levels, normalized to GUSB mRNA, of the selected transcripts analyzed by RT-qPCR upon depletion of GIGYF2 or control conditions. Mean values are shown with SEM for three to four independent experiments. Bottom: GIGYF2 knock-down was confirmed by western blotting. *P-value<0.01; **p<0.05; n.s., not significant. (D) Top: Enrichment of the selected transcripts in GIGYF2 immunoprecipitates analyzed by RT-qPCR. RNA samples were purified from input cell lysates and immuno-precipitation samples performed with an antibody against GIGYF2, or a negative control antibody (Control IgG). Transcripts levels in the IP were normalized to the input RNA fraction and enrichment in the GIGYF2 IP over the control IP was calculated. As a negative control, Log2 enrichment obtained for GUSB mRNA was set to 0. Mean values are shown with SEM. Bottom: Efficiency of the GIGYF2 Immunoprecipitation was analyzed by Western blotting.

### SVOPL and COL8A1 are repressed independently of m4EHP

Previously, a model was proposed in which GIGYF2 would act as a bridging protein between the cap binding protein m4EHP and RBPs to promote repression of specific transcripts (12). By contrast, the tethering assay data suggest an alternative model in which GIGYF2 recruit the CCR4/NOT complex to its bound mRNAs and promote their repression independently of m4EHP. To test which mechanism holds true for our identified targets, the variant of GIGYF2 unable to bind to m4EHP (mut GIGYF2) was overexpressed in HeLa cells and the outcome on SVOPL and COL8A1 expression levels analyzed by RT-qPCR (Figure 8A). When compared to conditions in which WT GIGYF2 was overexpressed we found that both SVOPL and COL8A1 mRNA were down regulated to a similar extent, demonstrating that m4EHP binding to GIGYF2 does not play a role for the repression of these transcripts (Figure 8B).

## DISCUSSION

Murine GIGYF2 was initially identified as a binding partner for the adaptor protein Grb10 that binds to phosphorylated insulin/IGF receptors and modulates their signaling (8). In mouse, attenuated expression levels of GIGYF2 leads to age-related motor dysfunction associated with hallmarks of neuronal disorders (11). It has been proposed that this phenotype is due to defects in insulin/IGF signaling caused by insufficient levels of cellular GIGYF2 (11,36). Recently, GIGYF2 was shown to be part of a translation repression complex when associated with the alternative cap-binding protein m4EHP (12). Here we have uncovered a novel feature of GIGYF2 as a direct RNA-binding protein that represses its targets independently of m4EHP through the recruitment of the CCR4/NOT complex.

### GIGYF2 is an RBP

GIGYF2 was identified as a potential RBP from mRNA interactome datasets in HEK-293 (1), mES (3), HepG2 (26) but not in HeLa cells (2). Since not all proteins identified in mRNA interactomes are *bona fide* RBPs (37), we have directly addressed whether GIGYF2 is an RBP in HeLa cells using a fluorescence-based ratiometric assay that measures the amount of polyadenylated RNA associated with a GFP-tagged protein(21). With this assay, GIGYF2 was clearly identified as an RBP. Interestingly, GIGYF2 does not display any predicted RNA binding domain and is thus one of the so-called “enigmRBP” with no clear function assigned to the RNA binding activity (26). We note that GIGYF1, the closest homolog of GIGYF2 in mammals with ca. 50% identity, was not found in the mRNA interactomes datasets and thus may not bind RNA. Interestingly, when compared to GIGYF1, GIGYF2 presents many more copies of tripeptide repeat motifs that are generally enriched in RBPs (26). This may guide further studies aimed at identifying the RNA binding domain of GIGYF2.

### GIGYF2 is a repressor of mRNAs through the recruitment of the CCR4/NOT complex

Using RNA tethering assays, we have shown that GIGYF2 represses bound mRNA by stimulating their decay. Furthermore, we have defined two distinct domains of GIGYF2 that are responsible for its silencing activity: the GYF and MED domains. We deciphered the mode of action of GIGYF2 - mediated repression by showing that it depends on the deadenylation activity of the CCR4/NOT complex that GIGYF2 recruits to its targets through distinct domains. Surprisingly, depletion of CNOT1 did not affect GIGYF2-mediated silencing. Altogether this suggests that binding of GIGYF2 to the CCR4/NOT complex involves multiple interfaces that engage several subunits of the complex.

### Multiple modes of repression for GIGYF2

GIGYF2 has been described as part of a translation inhibition complex through its binding to the cap binding protein m4EHP (12). In this context it was proposed that GIGYF2 might act as a bridging protein between RBPs and m4EHP to create a non-productive cap-binding complex. Such a mechanism appears to apply to TTP that binds m4EHP through GIGYF2 to enhance repression of its targets (13). Interestingly, the Izzauralde group recently reported that binding to GIGYF2 rather than cap-binding is essential for repression when m4EHP is artificially tethered to the 3’UTR of a reporter mRNA (38). However, the assembly of an intact m4EHP/GIGYF2 complex that binds to the cap proved to be required for full repression of a reporter harboring TTP recognition sites as a model for an endogenous target of m4EHP/GIGYF2 (38). By contrast, we propose an alternative mechanism of mRNA repression where GIGYF2 may directly bind to its target mRNAs independently of m4EHP. In this mode of repression, GIGYF2 primarily promotes the destabilization of its targets through the deadenylation activity of the CCR4/NOT complex. Importantly we have identified endogenous mRNAs that are repressed by GIGYF2 independently of m4EHP. We thus propose that GIGYF2 has a dual mode of action to repress target mRNAs. The use of multiple pathways by RBPs to achieve mRNA repression seems to be a more general feature than previously anticipated. For example, similar to GIGYF2, Smaug can repress translation through its interaction with the eIF4E-binding protein Cup (39), and stimulate mRNA decay through the recruitment of the CCR4/NOT complex (40). Whether GIGYF2 uses both modes of repression, sequentially or simultaneously, on the same transcripts is currently unknown. Alternatively, analog to TRIM71 in *C. elegans* (41), GIGYF2 may have a set of targets that are translationally repressed and another set that are degraded. How GIGYF2 switch from one mechanism to the other may then depend on the nature or the location (41) of the *cis*-regulatory elements displayed by its targets. The identification of the features recognized by GIGYF2 and the transcriptome-wide identification of its targets will help shed light on this issue.

### Interplay between GIGYF2 and miRISC

Previously we have shown that GW182 proteins directly and functionally interact with GIGYF2 (7). Indeed, GIGYF2 positively regulates miRNA-mediated translation repression but has no apparent effect on miRNA-mediated mRNA decay. The CCR4/NOT complex is also directly recruited by GW182 and stimulates mRNA deadenylation and decay (34,42,43). Based on these observations, we propose two possible models for the interplay between GIGYF2 and miRISC. In the first one, binding of GIGYF2 to GW182 or the CCR4/NOT complex is mutually exclusive. In that model, upon binding to an mRNA target, the Ago/GW182 complex would first recruit GIGYF2 and use the GIGYF2/m4EHP axis to repress translation. Thereafter the complex would remodel leading to recruitment of the CCR4/NOT complex and the release of GIGYF2/m4EHP. This remodeling would reflect the switch from translational repression to mainly stimulation of mRNA deadenylation and decay. As both GW182 and GIGYF2 were shown to directly bind to the CNOT9 (27,44,45), this subunit might be part of an overlapping interface. In the second model, GIGYF2 and GW182 can simultaneously bind to the CCR4/NOT complex. In that case the presence or absence of GIGYF2 may influence the importance of the translational repression component of miRNA-mediated silencing through the GIGYF2-mediated recruitment of m4EHP. This model would be in agreement with previous data suggesting that the CCR4/NOT complex is required for both miRNA-mediated translational repression and mRNA decay(46). Not integrated in these two models is the recent observation that m4EHP can also associate with CCR4/NOT through interactions with the proteins 4E-T and DDX6, and thereby regulate miRNA-mediated repression (47), suggesting that several ways of recruiting m4EHP to the miRISC co-exist. Studies aiming at defining the exact binding interfaces between GIGYF2 and the CCR4/NOT complex complemented with *in vitro* binding and functional assays should help deciding which of the two models better describes the interplay between GIGYF2 and the miRISC.

## Supplementary Data

Supplementary Data are available.

## Acknowledgement

The authors thanks Dr. Kai Schönig (Central Institute of Mental Health, Mannheim, Germany) for the kind gift of the HeLa-11ht cell line. Dr. Marina Chekulaeva (Max Delbrück Center, Berlin, Germany), Dr. Christian Freund (Freie Universität Berlin, Germany), Dr. Elisa Izzauralde (Max Planck Institute for evolution biology, Tübingen, Germany), Dr. Witold Filipowicz (FMI, Basel, Switzerland), Dr. Ann-Bin Shyu (University of Texas at Houston, USA), Dr. Sebastiaan Winkler (University of Nottingham, UK) for kindly sharing plasmids. We also thank Anne-Marie Alleaume and Dr. Matthias Hentze (EMBL, Heidelberg, Germany) for their help in establishing the RNA-binding protein assay as well as Jelena Pistolic and Dr. Vladimir Benes at the EMBL Genecore facility for the microarray analysis.

